# Extending mathematical frameworks to investigate neuronal dynamics in the presence of microglial ensheathment

**DOI:** 10.1101/2024.10.07.617090

**Authors:** Nellie Garcia, Gregory Handy

## Abstract

Recent experimental evidence has shown that glial cells, including microglia and astrocytes, can ensheathe synapses, positioning them to disrupt neurotransmitter flow between pre- and post-synaptic terminals. This study extends micro- and network-scale theoretical models to explore how varying degrees of synaptic ensheathment affect synaptic communication and network dynamics. Consistent with previous studies, our microscale model shows that ensheathment accelerates synaptic transmission while reducing its strength and reliability, with the potential to effectively switch off synaptic connections. Building on these findings, we integrate an “effective” glial cell model into a large-scale neuronal network. Specifically, we analyze a network with highly heterogeneous synaptic strengths and time constants, where glial proximity parametrizes synaptic parameters. Unlike previous models that assumed normal parameter distributions, our model uses parameters drawn from distinct distributions. This framework is applied to large networks of exponential integrate-and-fire neurons, extending linear response theory to analyze not only firing rate distributions but also noise correlations across the network. Despite the significant heterogeneity in the system, a mean-field approximation accurately captures network statistics. We demonstrate the utility of our model by reproducing experimental findings, showing that microglial ensheathment leads to post-anesthesia hyperactivity in excitatory neurons of mice. Furthermore, we explore how glial ensheathment may be used in the visual cortex to target specific neuronal subclasses, tuning higher-order network statistics.

## 1 Introduction

Recent advancements in experimental techniques and approaches in neuroscience have expanded the field beyond the traditional excitatory-inhibitory framework, revealing that various neuronal subtypes and glial cells significantly influence cortical dynamics. Instead of being a homogeneous class, 80% of interneurons are divided into three subtypes: parvalbumin (PV)-, somatostatin (SOM)-, and vasointestinal peptide (VIP)-expressing neurons, each with distinct properties (Pfeffer et al., 2013; Jiang et al., 2015; Karnani et al., 2016; Tremblay et al., 2016). Additionally, astrocytes and microglia, types of glial cells, have been found to closely ensheathe synapses, positioning them to fine-tune synaptic properties (Ventura and Harris, 1999; Chever et al., 2016; Haruwaka et al., 2024). These findings highlight the need for theoretical frameworks that incorporate cellular heterogeneity and offer deeper insights into how the brain utilizes this diversity in cortical computations.

Glial cells are thought to engage in bidirectional communication with neurons through several pathways. Experimental evidence supports their involvement in neurotransmitter clearance from the synaptic cleft, alterations of extracellular ion concentrations, the release of neuroactive substances via gliotransmission, and synaptic pruning (Tzingounis and Wadiche, 2007; Wake et al., 2009; Paolicelli et al., 2011; Covelo and Araque, 2018). Existing computational models have explored some of these functions, focusing on the concept of the “tripartite synapse,” which includes the pre- and post-synaptic terminals, as well as a neighboring astrocyte (Araque et al., 1999). These models, which primarily depend on gliotransmission, have demonstrated how astrocytes modulate network behavior, including influencing long-term potentiation/depression and modulating thresholds for focal seizure generation (Reato et al., 2012; Amiri et al., 2012; De Pittà and Brunel, 2016).

However, these models assume that the strength of astrocytic influence does not vary from synapse to synapse, despite evidence suggesting that astrocyte proximity to synapses, referred to here as ensheathment strength, is heterogeneous, varying across brain regions and disease states such as epilepsy and Alzheimer’s disease (Lippman et al., 2008; Coulter and Steinhauser, 2015; Matias et al., 2019; Price et al., 2021). Furthermore, recent experimental work by Haruwaka et al. (2024) demonstrated that changes in microglial ensheathment alone were sufficient to induce changes in network firing rates by shielding inhibitory synapses and disrupting neurotransmitter flow. This highlights a more subtle effect that both astrocytes and microglia can have on a network—an effect that does not rely on the sometimes controversial mechanism of gliotransmission (Nedergaard and Verkhratsky, 2012; Fujita et al., 2014; Haydon and Nedergaard, 2015). A recent model by Handy and Borisyuk (2023) investigated this effect with a detailed microscale model of synapses and derived an “effective” glial ensheathment framework that could be incorporated into large-scale neural networks. This computational study suggested that changes in synaptic strength and time course brought on by glial ensheathment could induce shifts in neural network synchrony. However, this work has a few shortcomings. First, the level of heterogeneity remained constrained, as the authors considered a binary level of ensheathment (each synapse was either ensheathed or not), while experiments show a distribution of ensheathment levels (Haruwaka et al., 2024). Second, their network model consisted of only two populations—excitatory and inhibitory neurons—and they primarily focused on the ensheathment of outgoing excitatory connections. Finally, it was left as an open question whether this heterogeneity could be accurately accounted for in a simplified mean-field model.

In this work, we aim to overcome these shortcomings by improving upon the “effective” glial model and incorporating it into a model of mouse primary visual cortex (V1), which accounts for the additional interneuron subclasses of PV, SST, and VIP. Recent experimental and modeling work has suggested that recurrent neuronal networks utilize these different inhibitory neurons to robustly perform complex cortical computations, such as tuning local and global oscillatory dynamics, recovering neuronal gain after injury, and shaping responses to visual stimuli during locomotion (Veit et al., 2023; Kumar et al., 2023; Dipoppa et al., 2018). This has led to the conjecture that there is a strong division of labor between interneurons, with different interneuron subtypes serving distinct roles in network activity and computation (Bos et al., 2020a). With this hypothesis in mind, it remains an open question whether glial ensheathment can modulate network dynamics in a similar fashion by targeting specific interneuron subtypes, motivating our choice of network configuration. Specifically, we will examine whether glial ensheathment can modulate the strength and synchrony of visually induced gamma oscillations, which are rhythms thought to promote the contextual synthesis of visual percepts (Fries, 2009).

In addition to constructing a spiking neuronal network that accounts for the heterogeneity induced by glial cells ensheathing specific synapses, we seek to extend linear response theory and develop a mean-field approximation that captures these results. This theory has been leveraged previously to investigate how neuronal firing rates and correlations respond to stochastic fluctuations (e.g., noisy inputs and synaptic connections) in their environment (Doiron et al., 2004; Lindner et al., 2005; Trousdale et al., 2012; Ocker et al., 2015). Recently, Veit et al. (2023) applied such theory to a network to investigate how these interneurons sub tune the strength of gamma rhythms. After extending this theory to include glial ensheathment, we will use it to perform an expansive parameter sweep, exploring the range of effects that glial ensheathment can have on network dynamics.

The paper proceeds with the following structure. First, we detail the modeling frameworks (Section 2), including the microscale model of glial ensheathment and the network model. In Section 2.1, we refine the “effective” glial ensheathment model presented in Handy and Borisyuk (2023) to fit a more realistic synaptic kernel and achieve greater accuracy across a range of ensheathment strengths. Section 2.2 then details the network model, generalizing previous work by accounting for multiple inhibitory subtypes and varying levels of glial ensheathment in a spiking network and its corresponding mean-field approximation. We then present key results in Section 3, where we replicate the experimental findings of Haruwaka et al. (2024), demonstrating that ensheathment of inhibitory synapses leads to hyperexcitability. We go beyond those results to explore how glial ensheathment affects correlations, particularly in modulating the strength and synchrony of gamma rhythms. After validating the match between spiking simulations and the mean-field approximation, we use our theory to conduct a large parameter sweep, comparing the effects of ensheathing PV neurons with those of SST neurons. Finally, in Section 4, we discuss these key results, model limitations, and future directions.

## 2 Models and Linear Response Theory

### 2.1 Microscale model of glial ensheathment

Haruwaka et al. (2024) observed that microglia ensheathed individual inhibitory synapses to varying degrees, including no contact, low contact, enwrapment, and complete engulfment, and that the overall level of ensheathment changed before and after anesthesia induced by the administration of isoflurane. Imaging data suggested that this ensheathment “shielded” pre-synaptic terminals, preventing neurotransmitters from diffusing across the synaptic cleft to the adjacent post-synaptic terminal. In this work, we utilize the diffusion with recharging traps (DiRT) process (Handy et al., 2018, 2019; Handy and Lawley, 2021) to model this shielding process at varying levels of glial cell ensheathment. We then generalize and refine the “effective” glial model of ensheathment developed in Handy and Borisyuk (2023) to more accurately capture the simulation results from this microscale model.

#### 2.1.1 Diffusion with recharging traps in an idealized synaptic cleft

We start by considering *N*_NT_ neurotransmitters diffusing within an idealized synapse, governed by the equation

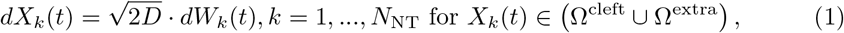

where *X*_*k*_(*t*) denotes the location of the neurotransmitter, *W*_*k*_(*t*) represents independent Wiener processes, and *D* is the diffusion coefficient. The synaptic cleft is modeled as a two-dimensional domain,

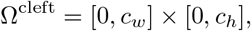

where *c*_*w*_ is the width and *c*_*h*_ is the height of the cleft. The boundary of the ensheathing glial cell is considered to be perfectly absorbing and is defined as

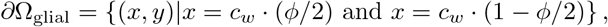

where *ϕ* ∈ [−1, 1] denotes the fraction of the synaptic cleft obstructed by glial protrusion. Note that *ϕ* can be negative (i.e., there exists space between the glial cell and the synaptic cleft), in which case the domain outside of the cleft is referred to as the extracellular space (Ω^extra^). Fig 1 illustrates this domain for various levels of glial ensheathment.

**Fig 1.**
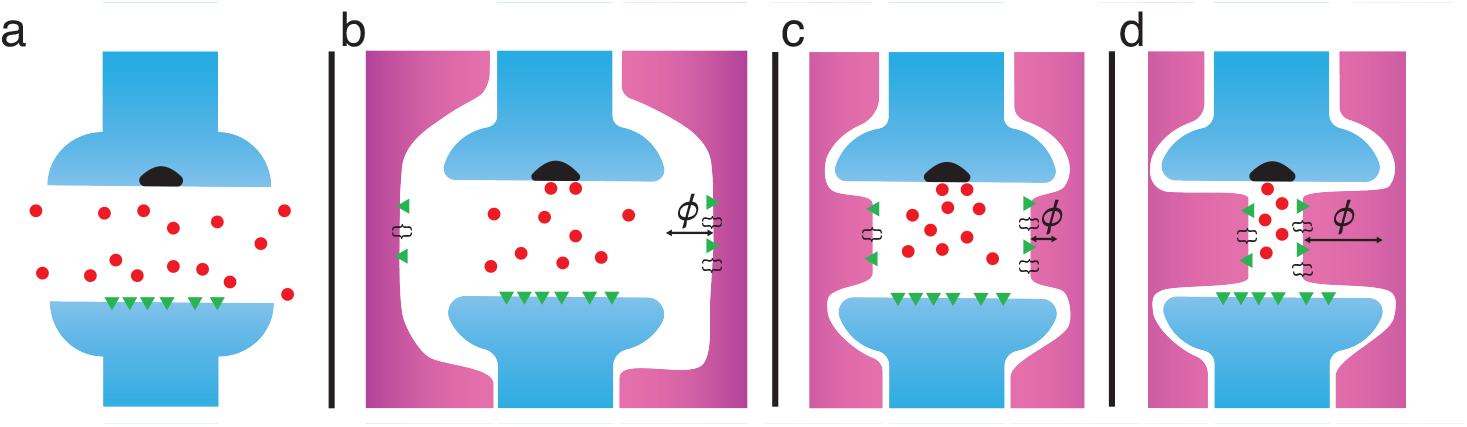
Schematics of the synaptic cleft, consisting of the pre- and post-synaptic terminals (blue) with different levels of glia (magenta) ensheathment. Neurotransmitters diffusing with the domain are denoted as red circles. a: no contact, b: low contact, c: high contact/enwrapped, and d: engulfed

Along the postsynaptic density, *N*_rec_ partially absorbing postsynaptic receptors of equal size are placed according to

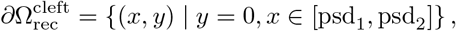

with the probability of a particle being absorbed upon making contact given by

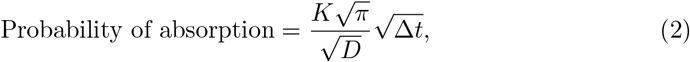

where Δ*t* is the time step of the diffusion model and *K* is the absorption rate. After a successful absorption, the receptor becomes activated and switches to a transitory refractory state, during which it is unable to bind additional molecules. The time spent in this refractory state is modeled as exponentially distributed, with a mean *τ*_*r*_ *>* 0. Once the receptor exits the refractory state, it returns to being partially absorbing. The remaining boundaries (i.e., the remaining portions of the pre- and post-synaptic terminals) are considered reflecting.

#### 2.1.2 Numerical details for the DiRT Simulations

We use the Euler-Maruyama method (Kloeden and Platen, 1992) for simulating Eq 1, implemented in a combination of C and MATLAB (2023). If a molecule hits an available receptor, it binds to the receptor with the probability given by Eq 2. If a receptor is in the refractory state, any neurotransmitter that hits it is reflected back into the domain. A bound neurotransmitter is removed from the system after an exponentially distributed time with rate *τ*_*r*_. All parameters corresponding to this ensheathment simulation can be found in Table 1.

**Table 1.** Default parameter value for the DiRT model (arbitrary units).

| Parameter | Default value | Description |
| --- | --- | --- |
| $N_{\text{NT}}$ | 1000 | number of neurotransmitters released |
| $N_{\text{rec}}$ | 50 | number of postsynaptic receptors |
| $\tau_r$ | 0.1 | mean receptor recharge time |
| $K$ | 1 | absorption rate |
| $D$ | 1 | diffusion coefficient |
| $c_w$ | 1 | cleft width |
| $c_h$ | 0.1 | cleft height |
| $[\text{psd}_1, \text{psd}_2]$ | [0.25 0.75] | postsynaptic density location |
| $\phi$ | -1 to 0.9 | fraction of the cleft blocked by protruding glial cell |
| $\Delta t$ | $1 \cdot 10^{-6}$ | diffusion time step |

#### 2.1.3 DiRT simulations and “effective” glial ensheathment model

Simulations of this stochastic process demonstrate how the time course of the number of active receptors is influenced by glial ensheathment via changes in the protrusion parameter *ϕ* (Fig 2a). We find that as *ϕ*→ 1, both the maximum number of activated receptors and the time course of this activation significantly decrease. To further quantify this change, we fit the curves to *α*-functions, a standard model of synaptic interactions (Trousdale et al., 2012),

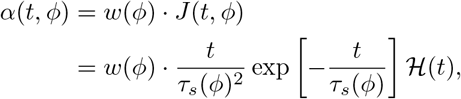

where we have normalized the *α*-function so that

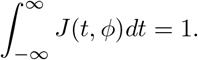

**Fig 2.**
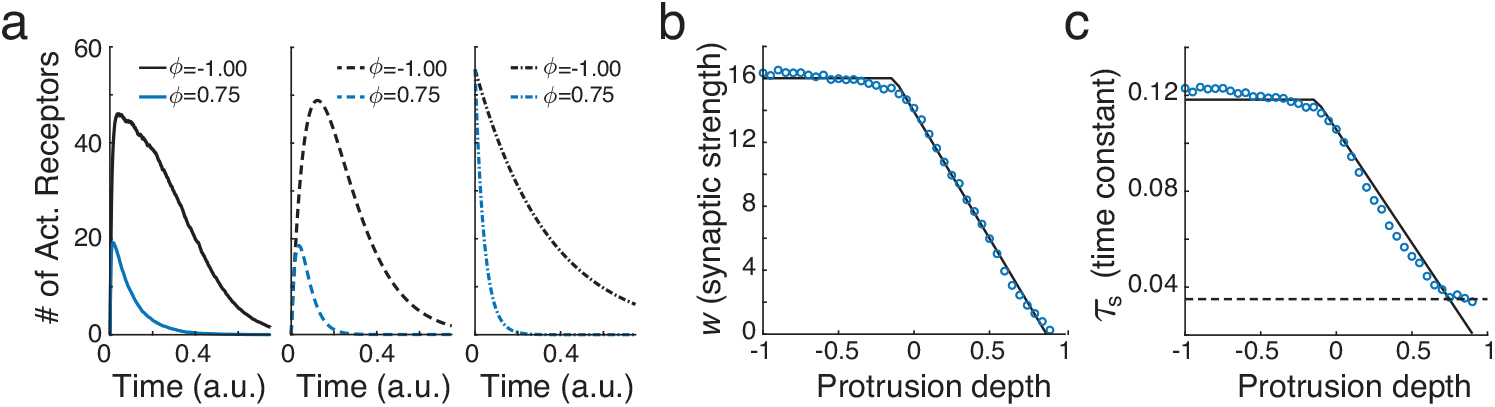
a: Simulation (left panel) and fitting results (middle panel: *α*-function fits; right panel: exponential fit from the model used in Handy and Borisyuk (2023)) of neurotranmitters diffusing in the synaptic clef with glial ensheathment for *ϕ* = *−*1 (black) and *ϕ* = 0.75 (blue). b: Fitted synaptic strengths, *w*(*ϕ*) for different protrusion amounts (blue circles) and the fitted piecewise function from Eq 3 (black). c: Same as b, except for the synaptic time constants *τ*_*s*_(*ϕ*). The black dash line indicates the minimum time constant reached, 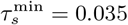

In this formulation, *w*(*ϕ*) and *τ*_*s*_(*ϕ*) correspond to the synaptic strength and synaptic time constant, respectively, for different levels of *ϕ*. Fig 2a (middle panel) illustrates that this function provides a reasonable fit for *ϕ* = −1 and *ϕ* = 0.75 when compared to the DiRT simulations (*R*^2^ = 0.951 and 0.929 for these two values of *ϕ*, respectively).

Varying *ϕ* ∈ [−1, 0.9], we observe that *w*(*ϕ*) and *τ*_*s*_(*ϕ*) follow a piecewise linear pattern, qualitatively similar to that found in Handy and Borisyuk (2023) (Fig 2c and 2d, blue circles). Specifically, for negative *ϕ, w*(*ϕ*) and *τ*_*s*_(*ϕ*) remain constant, indicating that the neighboring glial cell has no effect on synaptic transmission. Both values then decrease linearly as *ϕ* increases towards 1. Assuming this general form, we estimate the transition point by fitting a piecewise linear function:

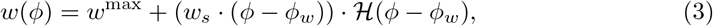

to the simulated data points (Fig 2c and 2d, black line). We find *w*^max^ = 15.747, *w*_*s*_ = −15.586, and *ϕ*_*w*_ =− 0.124. Interestingly, this suggests that glial cells can influence synaptic transmission by simply being in the *proximity* of a synapse (i.e., direct contact is not necessary). Furthermore, *w*(*ϕ*) → 0 as *ϕ* →1. The fit for *τ*_*s*_(*ϕ*) is similar, except that 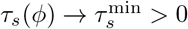as *ϕ*→ 1.

Performing such detailed diffusion simulations of neurotransmitters is infeasible in the context of a large-scale spiking neuronal network, but Eq 3 suggests an effective glial ensheathment model that can be readily implemented to account for changes in synaptic strength and time constant. Specifically, let *w*_*ab*_ be the default (maximal) synaptic strength from neuron population *b* to *a, s*_*en*_ ∈ [0, 1] denote the degree of ensheathment (i.e., *s* = 0 indicates no ensheathment, and *s* = 1 indicates full engulfment), and **1**_*ij*_ be an indicator function that equals 1 if the *ij* synapses is ensheathed. Approximating *w*_*s*_ ≈ −*w*_*ab*_ and substituting *s*_*en*_ = *ϕ* − *ϕ*_*w*_ into Eq 3 gives:

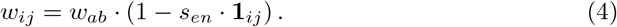

The equation for 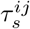 is similar, except we account for its minimal value, 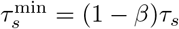 for *β* ∈ (0, 1), as *s*_*en*_ → 1:

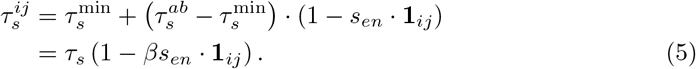

This model of an “effective” glial cell can be readily implemented into a large-scale network, as discussed below. It is similar to the results found in Handy and Borisyuk (2023), but differs in two ways. Here, we consider *α*-functions for the synaptic interactions and explicitly fit the synaptic strength and time constants to the simulation results. In the previous work, they considered exponential synapses and made phenomenological connections to parameter values. Furthermore, by accounting for 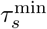, our model allows the maximal height of the synaptic interactions to vary, a feature that was not achievable in the previous model. Fig 2a (right panel) show the best-fit lines using this previous framework. Overall, this model provides a more accurate description of how glial ensheathment makes synapses faster and weaker.

### 2.2 Details and connectivity of the spiking neuronal network

#### 2.2.1 Exponential integrate-and-fire network with glial ensheathment

We consider a network of *N* recurrently connected exponential integrate-and-fire (EIF) neurons with membrane potentials of the form

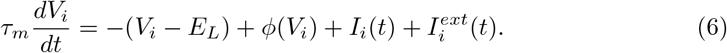

Each neuron *i* = 1, …, *N* belongs to one of three subclasses *a* = *e*_*x*_ (excitatory), *p*_*x*_ (PV), or *s*_*x*_ (SST) located at either the center (*x* = 1) or surround (*x* = 2). A neuron spikes when *V*_*i*_(*t*) ≥ *V*_*th*_, after which its value is reset to *V*_*re*_ and undergoes a refractory period of length *τ*_ref_. Here, *E*_*L*_ denotes the leak reversal potential and *ϕ*(*V*) represents the spike-generating current, which takes the form

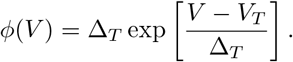

Synaptic interactions are modeled as

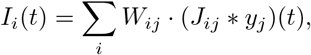

where the spike train from neuron *j* is the point process *y*_*j*_(*t*) = ∑_*k*_ *δ*(*t* −*t*_*j,k*_) and ∗ denotes convolution. Following the work from our microscale model in Secion 2.1.3, synaptic interactions are modeled by delayed *α*-functions

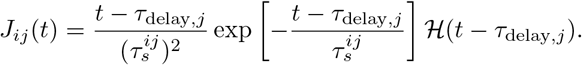

The synaptic weights are given by

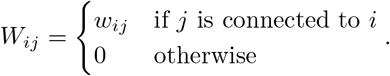

We implement the results of the microscale ensheathment model following Eqs 4 and 5, but generalizing to allow a network to have multiple levels of ensheathment strengths. Specifically, let 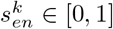 denote the *k* = 1, 2, …, *m* possible levels of ensheathment strengthens. Then, for a synaptic connection from neuron *j* to neuron *i*, we let 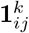 be an indicator function that equals 1 if the *ij* synapse is ensheathed at level *k* and zero otherwise. We can then write the synaptic strength and time constant for synapse *ij* as

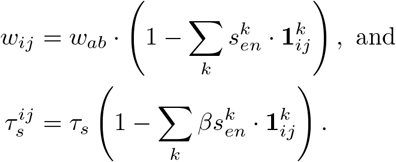

The last term of Eq 6, 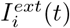, represents the external drive to the network. We consider both a fixed background input (driving all cells) and a fixed visual input stimulating the center and surround visual areas (driving only excitatory and PV cells). Following the work from Veit et al. (2023), VIP neurons are accounted for as a feedforward source of inhibition onto SST cells and play a key role in setting their baseline firing rate. Assuming all of these inputs arrive as Poisson spike trains, we make use of a diffusion approximation and Campbell’s theorem (Kingman, 1993) to write these inputs as

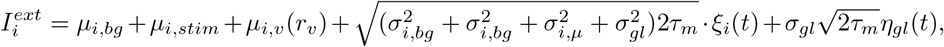

where *ξ*_*i*_(*t*) is a zero mean, delta-correlated ⟨*ξ*_*i*_(*t*), *ξ*_*i*_(*t*^*′*^)⟩ = *δ*(*t* − *t*^*′*^) Gaussian white noise term, and *μ*_*i,v*_ = (*w*_*i,v*_*N*_*v*_) · *r*_*v*_ and 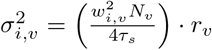. We denote the global noise process as *η*_*gl*_(*t*), which is primarily considered to be a zero mean, delta-correlated Gaussian white noise, though our corresponding linear response theory (Section 2.3) begins by considering it to be bandlimited. All default parameter values can be found in Table 2.

**Table 2.**
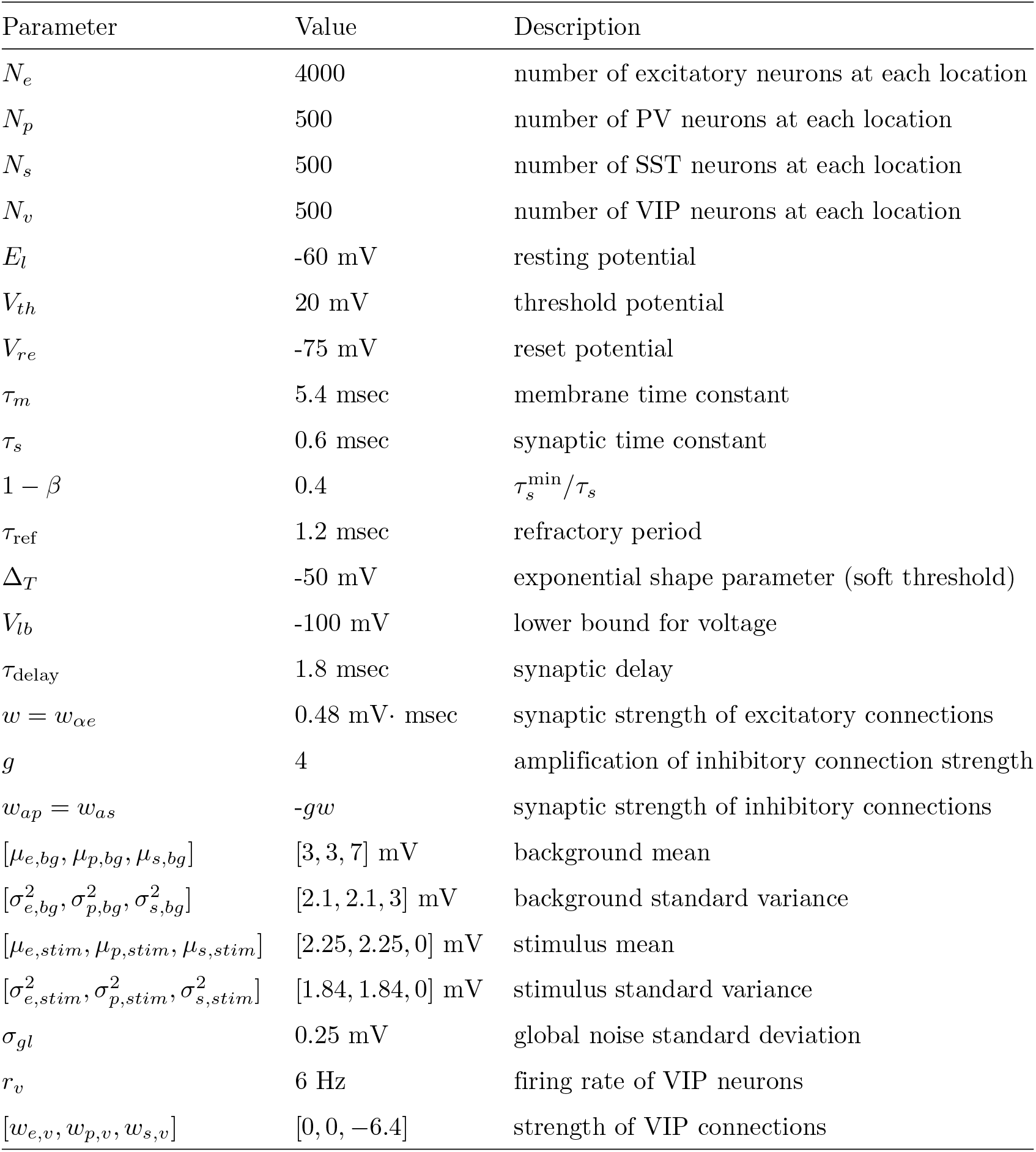
Default neuronal and network parameters. Changes to any parameter are indicated in the figure caption.

| Parameter | Value | Description |
| --- | --- | --- |
| $N_e$ | 4000 | number of excitatory neurons at each location |
| $N_p$ | 500 | number of PV neurons at each location |
| $N_s$ | 500 | number of SST neurons at each location |
| $N_v$ | 500 | number of VIP neurons at each location |
| $E_l$ | -60 mV | resting potential |
| $V_{th}$ | 20 mV | threshold potential |
| $V_{re}$ | -75 mV | reset potential |
| $\tau_m$ | 5.4 msec | membrane time constant |
| $\tau_s$ | 0.6 msec | synaptic time constant |
| $1 - \beta$ | 0.4 | $\tau_s^{\min}/\tau_s$ |
| $\tau_{ref}$ | 1.2 msec | refractory period |
| $\Delta_T$ | -50 mV | exponential shape parameter (soft threshold) |
| $V_{lb}$ | -100 mV | lower bound for voltage |
| $\tau_{delay}$ | 1.8 msec | synaptic delay |
| $w = w_{\alpha e}$ | 0.48 mV · msec | synaptic strength of excitatory connections |
| $g$ | 4 | amplification of inhibitory connection strength |
| $w_{ap} = w_{as}$ | $-gw$ | synaptic strength of inhibitory connections |
| $[\mu_{e,bg}, \mu_{p,bg}, \mu_{s,bg}]$ | [3, 3, 7] mV | background mean |
| $[\sigma_{e,bg}^2, \sigma_{p,bg}^2, \sigma_{s,bg}^2]$ | [2.1, 2.1, 3] mV | background standard variance |
| $[\mu_{e,stim}, \mu_{p,stim}, \mu_{s,stim}]$ | [2.25, 2.25, 0] mV | stimulus mean |
| $[\sigma_{e,stim}^2, \sigma_{p,stim}^2, \sigma_{s,stim}^2]$ | [1.84, 1.84, 0] mV | stimulus standard variance |
| $\sigma_{gl}$ | 0.25 mV | global noise standard deviation |
| $r_v$ | 6 Hz | firing rate of VIP neurons |
| $[w_{e,v}, w_{p,v}, w_{s,v}]$ | [0, 0, -6.4] | strength of VIP connections |

#### 2.2.2 Connectivity and ensheathment rules

For neurons located in the same location, we follow the connectivity rules outlined in previous experimental and modeling works (Pfeffer et al., 2013; Litwin-Kumar et al., 2016; Bos et al., 2020a; Veit et al., 2023), as illustrated in Fig 3a. Note that most populations share recurrent connections both across and within subpopulations, with the exception of SST neurons, which do not have recurrent connections to other SST cells or receive inputs from PV cells. We also consider long-range excitatory connections across our discrete spatial locations (see Fig 3b). These connections qualitatively align with the parameters used in Keller et al. (2020), which fitted a rate-based model to experimental data collected from L2/3 of a mouse presented with different visual stimuli.

**Fig 3.**
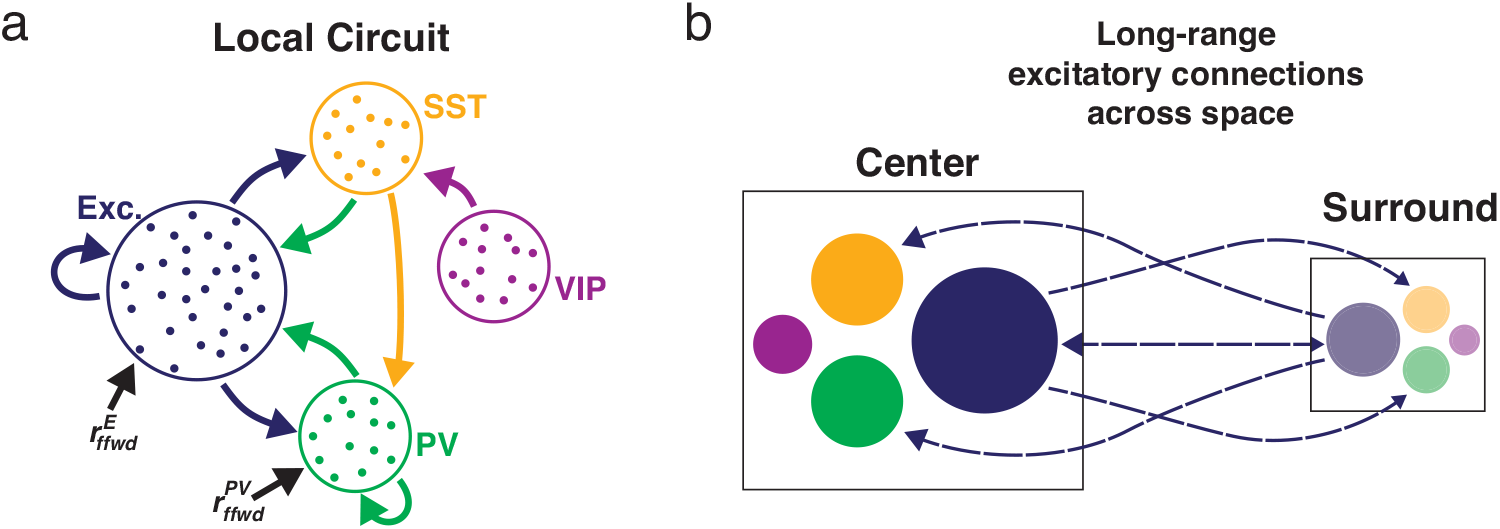
a: Network schematic showing the local connectivity rules across the neuronal populations (blue: excitatory, green: PV, yellow: SST, and magenta: VIP). b: Network schematic showing the long range excitatory connections between the center (solid colors) and surround (translucent colors) locations

To create the connectivity matrix *W*, each neuron *j* from population *b* is randomly connected (without replacement) to *p*_*ab*_*N*_*a*_ neurons from population *a*, resulting in a fixed out-degree network. The connection probability parameters are listed in Table 3. From these possible synapses, each one has a probability of 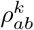 of being ensheathed with strength 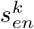. The parameters 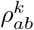 and 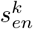 vary throughout this work, with their values clearly indicated within the figures. Taking inspiration from the experimental study by Haruwaka et al. (2024), we explore the effects of ensheathing inhibitory synapses.

**Table 3.**
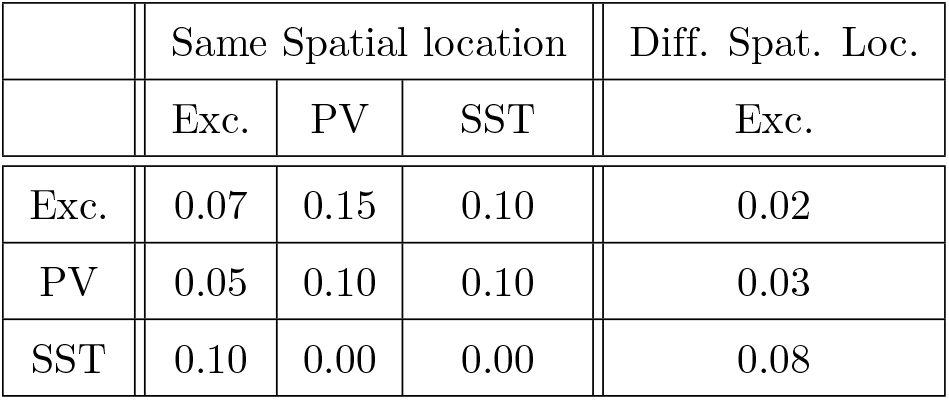
Probability of connections *p*_*ab*_ (from population *b* to *a*; columns presynaptic, rows postsynaptic). Changes to any parameter are indicated in the figure caption.

#### 2.2.3 Neuronal network simulation and network statistics

The spiking simulations were completed with Euler’s method using a time step of 0.025 msec for a total of 1· 10^6^ msec of simulation time. The output of cell *i* can be described by its output spike train

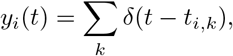

where the *t*_*i,k*_’s denote the times when the neuron’s membrane potential crosses the spike threshold *V*_*th*_. For the default parameter set without any glial ensheathment, we observed that the neurons in the network are spiking asynchronously, with low correlations within and across populations, as illustrated in the raster plot shown in Fig 4a (bottom). Fig 4b (top) shows the rolling firing rates for the two excitatory populations.

**Fig 4.**
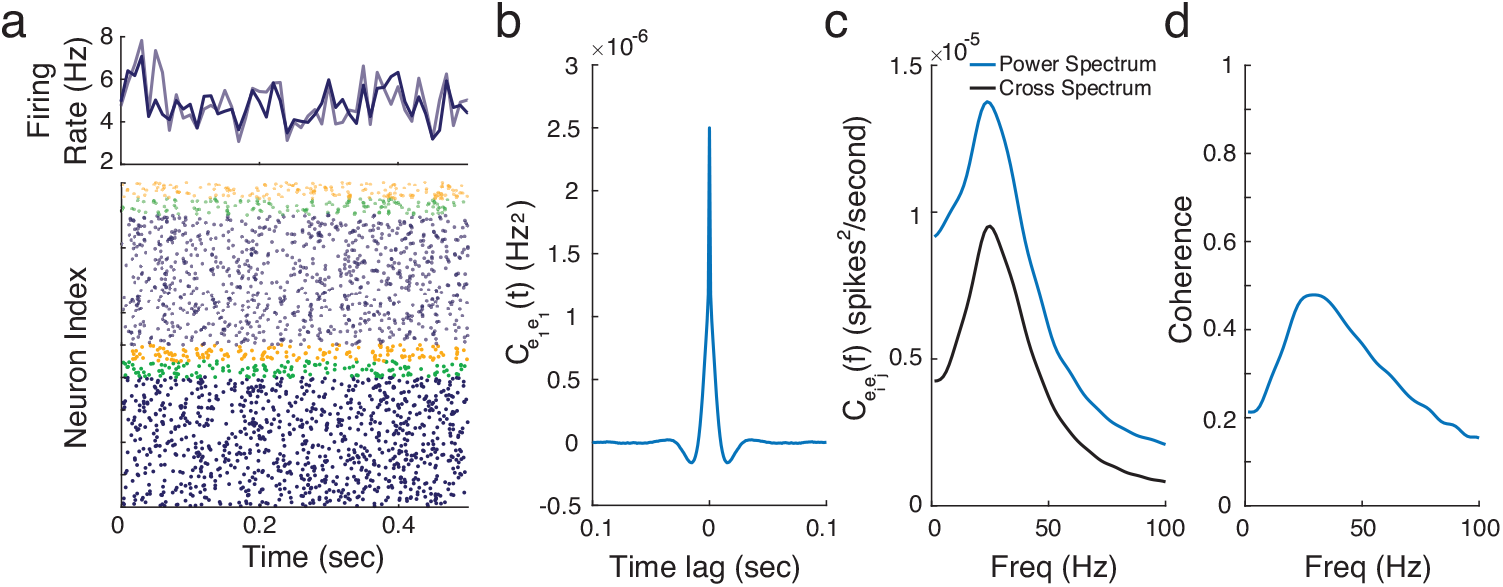
a: (bottom) Raster plot indicating the spike times of a sample of neurons (from the excitatory (blue), PV (green) and SST (yellow) populations at the center (solid) and surround (translucent), (top) Average firing rate (averaged over 0.01 second time windows) for the excitatory neurons in the two spatial locations. b: Auto-covariance function for the excitatory population for different time lags. c: The power spectrum for the excitatory population (blue) and cross-spectrum between excitatory populations at different spatial locations (black) across different frequencies. d: Coherence across frequencies between the two excitatory populations

In this work, we are interested in exploring how glial ensheathment can modulate average network statistics such as firing rates and correlations. We define the average firing rate of neuron *i* as *r*_*i*_ = ⟨*y*_*i*_(*t*)⟩ and the cross-correlation function of neurons *i* and *j* as *C*_*ij*_(*h*) = ⟨(*y*_*i*_(*t*) −*r*_*i*_)(*y*_*j*_(*t* + *h*) −*r*_*j*_)⟩, where ⟨·⟩ denotes the average across time. The average population firing rate is then

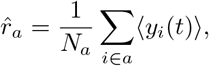

with the average population auto- and cross-covariance functions defined as

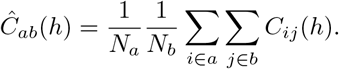

Numerically, these auto- and cross-correlation functions are estimated by binning the spike times over 0.001 second time windows, summing this count across subpopulations in different spatial locations, and then using MATLAB’s built-in xcorr() function with a maximum window length of 0.25 seconds. We can then take the Fourier transform of these functions to find the power spectrum, 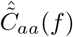, and cross-spectrum, 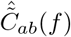. This allows us to examine the magnitude of these spectra at different frequencies. We will use these quantities to also estimate the coherence across spatial locations, which is defined as

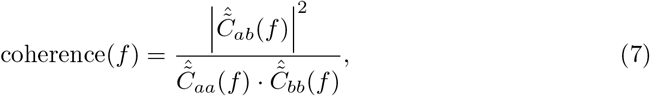

which is a normalized measure of synchrony, with values ranging from 0 (no synchrony) to 1 (high synchrony).

Fig 4b-4d shows the estimated auto-correlation, power, cross-spectrum, and coherence for the default network. For these parameter values, the network exhibits a strong gamma rhythm, evident by the bump in the spectrum occurring in the 20-50 Hz range, with moderate levels of coherence at these frequencies. We define the “gamma power” as the peak of the power spectrum within this frequency range, and the “gamma frequency” as the frequency at which this peak occurs. Additionally, we define “gamma coherence” as the coherence value at the corresponding gamma frequency.

### 2.3 Linear response theory

While the numerical simulations of the spiking network detailed in Section 2.2 can demonstrate how glial ensheathment of synapses modulates network dynamics, performing large parameter sweeps is impractical due to the computational cost. To address this, we develop a mean-field linear response theory that can be used for this purpose.

To begin, we first derive (Section 2.3.1) the average population steady-state firing rates and effective mean inputs using the self-consistency relationship:

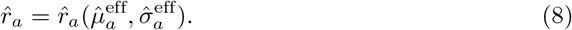

Next, we derive (Section 2.3.2) a formula for 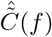, which is dependent on the linear response function, *Ã*_*a*_(*f*), and the power spectrum of the baseline spike trains, 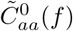. Both of these are defined in Section 2.3.2. With the results from Eq 8, we can apply the theory from Richardson (2007, 2008) to the following stochastic differential equation

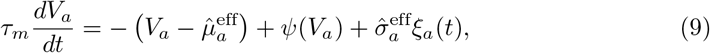

in order to numerically estimate these two functions, which can then be used in our calculation of 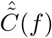.

#### 2.3.1 Average firing rate

We start by considering the steady state firing rates, which can be solved via the self-consistency relationship

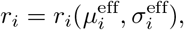

Where

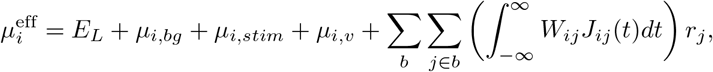

and

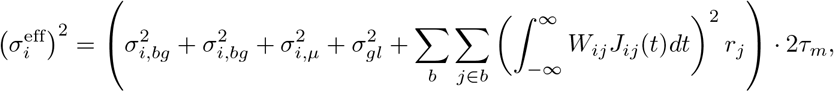

are the effective mean input and variance into neuron *i*, both of which depend on the firing rates *r*_*j*_. These forms follow from the a diffusion approximation and assumption that these recurrent inputs arrive as Poisson spike trains, as approximated in previous works (Trousdale et al., 2012; Veit et al., 2023). This relationship can be solved numerically using standard methods developed in Richardson (2007, 2008) for a nonlinear integrate-and-fire neuron and fixed point iteration.

In seeking a formula for the average firing rate across the populations, 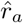, we begin by noting that the leak reversal potential, as well as the mean and variance of external inputs, are the same for all neurons in population *a*. Thus, the average effective input 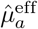 into population *a* can be written as

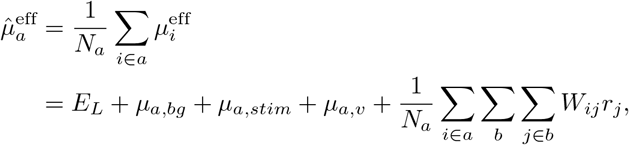

where we have also used the fact that the the integral of the synaptic interaction is 1 by design, even in the presence of glial ensheathment. We can approximate the last term in this sum as follows

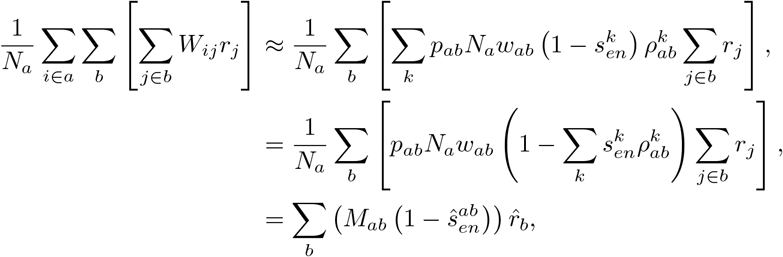

Where

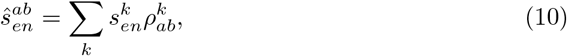

is the weighted average of the ensheathment strength from population *b* to *a*, and

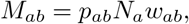

is the effective connectivity strength from population *b* to *a* in the absence of glial ensheathment. The first line follows from the fact that a neuron in population *b* makes on average 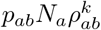 connections with an ensheathment strength of 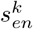 to a neuron in population *a*.

The calculation for the average variance proceeds in a similar fashion,

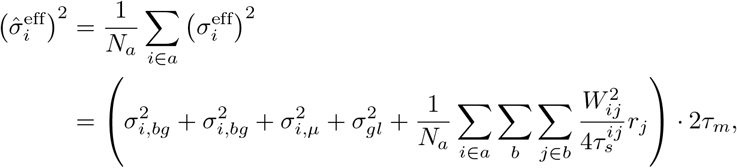

where we have calculated the integral of the square of the synaptic interaction term. The last term in the parentheses can be approximated similar to the steps above to find

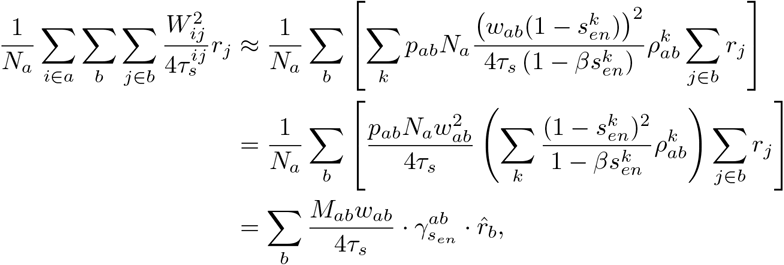

where

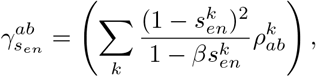

captures the effect ensheathment has on this higher-order correction term.

These calculations yield a self-consistency relationship for the average population firing rates

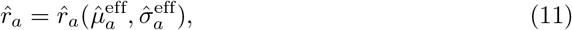

where

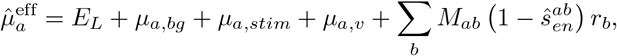

and

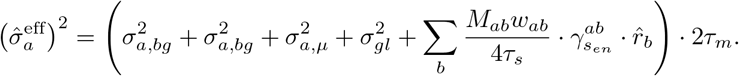

This result is written to explicitly highlight the dependence on ensheathment strength. When there is no ensheathment (i.e., 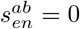 and 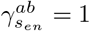), this self-consistency relationship simplifies to rely only on terms involving the traditional effective connectivity strength, as derived in previous mean-field analyses (e.g., Rosenbaum et al. (2017) and Bos et al. (2020b)).

#### 2.3.2 Average power- and cross-spectrums

We now seek to derive a formula for the power and cross-spectrum matrix, 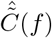. To do this, we follow previous work (Doiron et al., 2004; Lindner et al., 2005; Trousdale et al., 2012; Veit et al., 2023) and linearize each neuron’s spike train around a realization of the spiking output in the absence of recurrent connections, 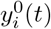. We begin by explicitly assuming that the synaptic connections, *W*_*ij*_, are weak and that the global noise process is band-limited. This allows us to approximate the spike response from neuron *i* in the Fourier domain as

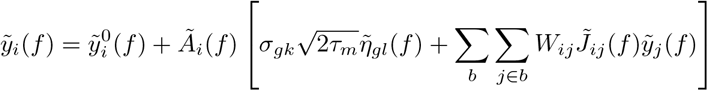

where 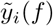 is the zero-mean Fourier transform of the spike train, 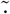 denotes the Fourier transform of the other quantities, and *Ã*_*i*_(*f*) is the linear response of the postsynaptic neuron (Gardiner, 2009). Averaging the spike response equation across population *a* then yields

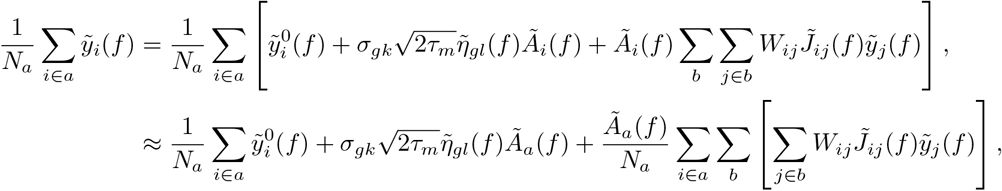

where we have made the approximation *Ã*_*a*_(*f*) ≈ *Ã*_*i*_(*f*). The third term in this sum can be simplified following the same reasoning as in Section 2.3.1,

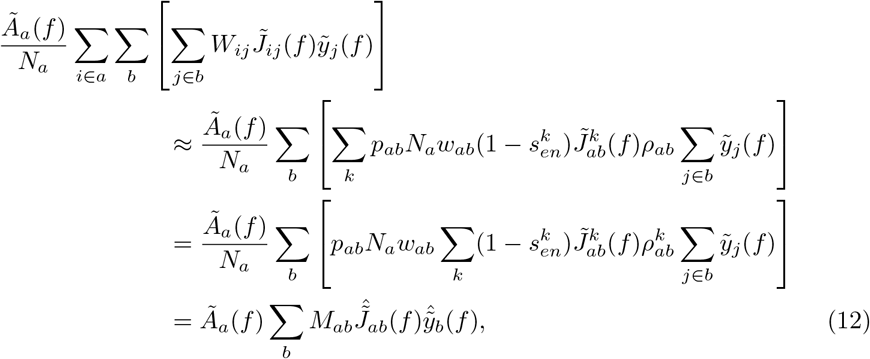

where

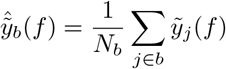

denotes the Fourier transform of the concatenated spike trains from each population,

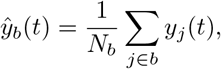

and

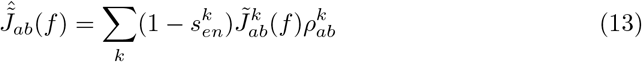

is a modified synaptic kernel that accounts for ensheathment strength. We note that in the case of a fixed in-degree network, *Ã*_*a*_(*f*) = *Ã*_*i*_(*f*) (i.e., all neurons have the same linear response function). Here, we consider a fixed out-degree network, which has the benefit of ensuring that a neuron in population *b* makes exactly *p*_*ab*_*N*_*a*_ connections to a neuron in population *a*, which appears in the derivation of Eq 12.

Using this simplified form, we can now write

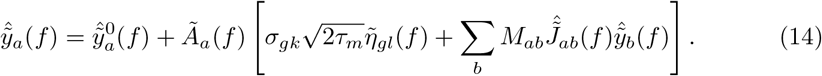

It is important to note that, unlike the final self-consistency relationship used to estimate average firing rates in Section 2.3.1, the effect of ensheathment strength here is incorporated into the modified synaptic kernel.

Eq 14 represents 6 ×6 linear system, the solution to which, 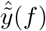, can be used to find the averaged power spectrum and cross-spectrum matrix. Aside from the use of our modified synaptic kernel, this calculation follows exactly as in Veit et al. (2023), and results in the following expression

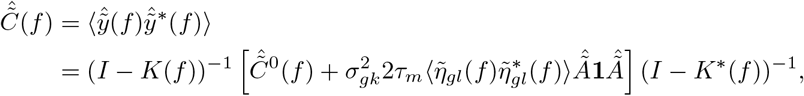

with

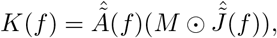

where 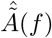 is a diagonal matrix of linear response functions, *M* is the matrix of effective connectivity strengths, 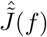 is the matrix of modified synaptic kernels, **1** is the matrix of all ones, and 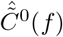 is a diagonal matrix with entries

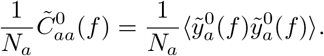

Similarly, following the work of Lindner et al. (2005) we can extend this to account for global noise that takes the form of Gaussian white noise (with infinite variance) and find

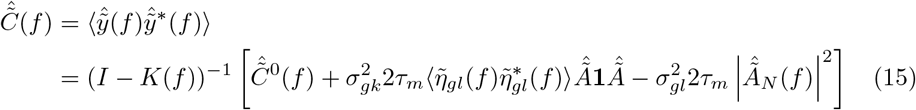

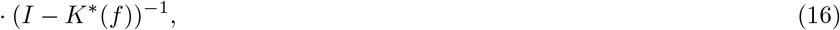

where 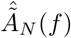 is a diagonal matrix with entries *Ã*_*a*_(*f*)*/N*_*a*_. In this work, we consider global noise with a magnitude small relative to the independent noise, leading to this correction term having a minimal effect, as it is orders of magnitude smaller than the other terms within the brackets.

While this theory allows for the probability of ensheathment to vary from population *b* to *a*, for the remainder of this work, we make the simplifying assumption that these values depend only on the pre-synaptic population and are the same for both the center and surround locations. As a result of this simplification, we can write 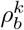 to represent the probability of a synapse from neuronal subclass *b* being ensheathed with strength 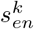.

#### 2.3.3 Comparison to naïve mean-field approach

It is worth noting that our self-consistency relationship (Eq 11), as well the average spike response from a neuron (Eq 14), vary from what one would find from a more naïve mean-field approach. Specifically, one could first average across the network to find the average level of ensheathment experienced across the populations (Eq 10), then update the synaptic weights and time constants to find

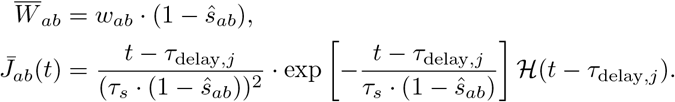

It is clear that these terms do not appear in our derivations, indicating that this naïve approach would lead to deviations in this approximation for the average firing rates, power spectrum and cross-spectrum.

## 3 Modeling results

### 3.1 Replication of experimental observations

With the spiking network and mean-field theory established, we first aim to replicate the recent results of Haruwaka et al. (2024). In that study, they observed that during emergence from anesthesia induced by the administration of isoflurane, neurons in layer 2/3 of the cortex became hyperactive, with a significant increase in firing rates. To explain these results, they tracked changes in microglia-neuronal interactions, specifically focusing on the number and strength of the connections between these two cell types. They found that the percentage of neurons with nearby microglia increased from 20% to nearly 80% from the awake to the anesthetized and post-anesthetized (i.e., “emergence”) states (Fig 5a). Interestingly, they also found that these interactions predominantly affected GABAergic synapses (i.e., inhibitory inputs). Using confocal microscopy, they classified these interactions into four subtypes based on surface contact: no contact, less than 50% surface coverage (low contact), more than 50% surface coverage (enwrapped), and complete engulfment. These levels also changed significantly across the three states, with an increase in enwrapped and engulfed synapses during and after isoflurane administration (Fig 5b). Finally, they confirmed through super-resolution and electron microscopy that the tips of microglial bulbous endings shielded PV pre-synaptic terminals by protruding into the cleft, consistent with our microscale model.

**Fig 5.**
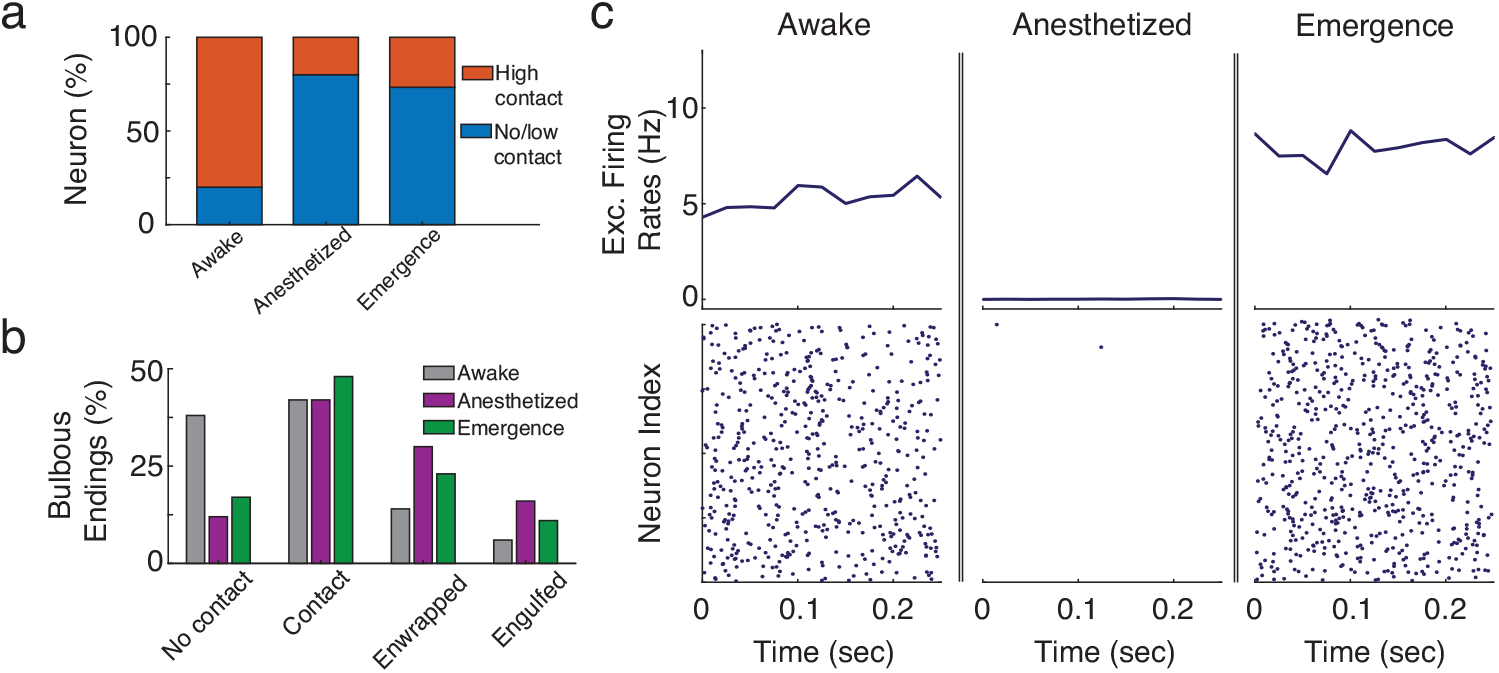
a: Modified panel from Haruwaka et al. (2024) showing that the number of neurons with multiple synapses with nearby microglia increases significantly after the administration of isoflurane, and remains after its removal (i.e., ‘emergence’). b: Modified panel from Haruwaka et al. (2024) showing how the coverage of GABAergic synapses change between the three states. c: Raster plots (bottom) and average firing rates (averaged over 0.25 second time windows) showing the simulation results across the three experimental states using ensheathment parameters found in Table 4. In addition, for the anesthetized state, the mean and variance of the feedforward input was set to [*μ*_*e*,ffwd_, *μ*_*p*,ffwd_, *μ*_*s*,ffwd_] = [0, 0, 0] mV, and 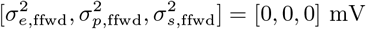

With these experimental findings as a reference, we ask whether our spiking model can replicate the observed increase in hyperexcitability from the awake to emergence state via changes in glial ensheathment alone. We consider four levels of ensheathment strength: 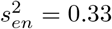, and 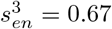. The probability of ensheathment for out-going inhibitory synapses (both PV and SST) was taken from the experimental data and can be found in Table 4. Meanwhile, all other synapses were placed in the no contact state. In addition to changing the ensheathment levels, the anesthetized state, which exhibits little to no spiking due to the inhibition of external driving forces, was modeled by removing the feedforward input.

**Table 4.**
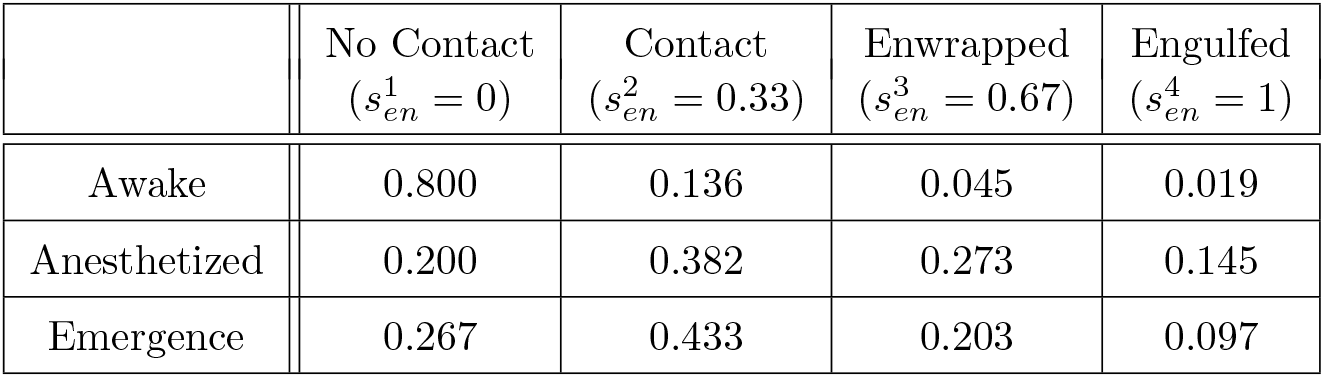
Probability of ensheathment for all outgoing inhibitory connections, *ρ*_*p*_ and *ρ*_*s*_, as used in Fig 5.

Fig 5c illustrates the results of the model with raster plots (top) and the rolling average firing rate (bottom) for the three different states. As expected, there is a significant drop in excitatory firing rates from the awake to the anesthetized state, which is not due to changes in glial ensheathment but rather the removal of feedforward inputs. Once the model enters the emergence state, where glial ensheathment levels remain elevated and the feedforward inputs are restored, the network enters a hyperactive state, showing a 43% increase in excitatory firing rates. These baseline results demonstrate that the “effective” glial ensheathment model derived in Section 2.1 can successfully replicate this key experimental result when implemented into a large-scale spiking model. We also report significant increases in firing rates across the PV and SST populations (Fig 6a). It is worth noting that although the percentages used in Table 4 are well estimated from the experimental data, the corresponding strength of ensheathment values are not. However, the qualitative behavior observed here is robust to these parameter choices.

**Fig 6.**
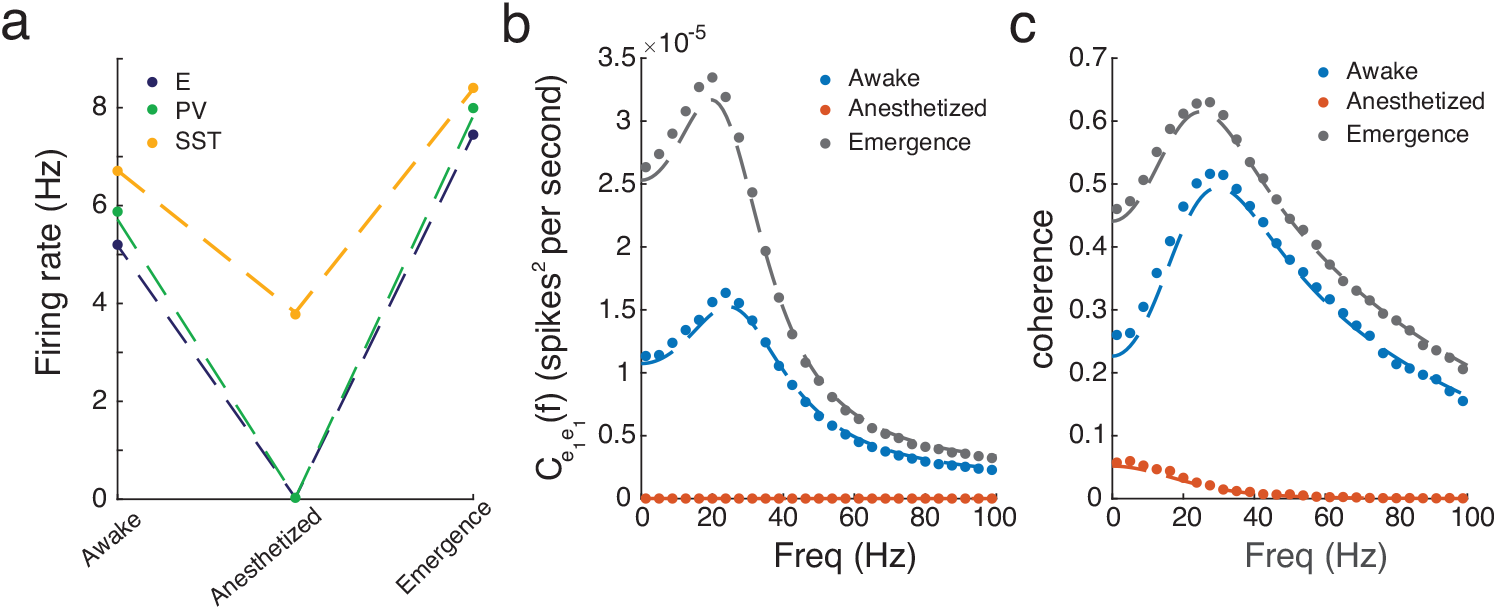
Results of the spiking model (dots) and mean-field approximation (dashed-line) for the a: firing rates, b: power spectrum, and c: coherence across the awake, anesthetized and emergence states using the ensheathment parameters found in Table 4. In addition, for the anesthetized state the mean and variance of the feedforward input was set to [*μ*_*e*,ffwd_, *μ*_*p*,ffwd_, *μ*_*s*,ffwd_] = [0, 0, 0] mV, and 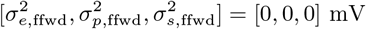

### 3.2 Higher ordered statistics and verification of linear response theory

The implications of this hyperactivity on cortical processes that depend on higher-order network statistics remain an open question. Here, we focus on network dynamics in V1, examining the strength and synchrony of gamma rhythms, which are thought to play a role in spatially integrating visual information. Following the steps in Section 2.2.3, we calculate the average power spectrum and cross-spectrum for the excitatory populations at both spatial locations. Fig 6b shows a clear increase in the power spectrum across all frequencies, with the largest increase occurring within the gamma frequency range. Similarly, Fig 6c shows a significant increase in the coherence (i.e., a normalized measure of synchrony; see Eq 7) of these gamma oscillations. Thus, in addition to shaping the firing rates of the network, our model predicts that glial ensheathment can lead to significant changes in synchronous activity. This is also a testable prediction. Veit et al. (2017, 2023) examined coherence across V1 in head-fixed mice viewing oriented gratings in the context of optogenetic stimulation of different interneuron subclasses. It would be interesting to replicate those experiments as the animal emerges from an anesthetized state to see if these increases in gamma power are observed.

With this parameter set and results in mind, we now turn to the linear response theory and the corresponding mean-field approximation derived in Section 2.3 to determine if similar results hold. We find impressive agreement for the firing rates, power spectrum, and coherence in both the awake and emergence states (Fig 6). Recall that the heterogeneity induced by glial ensheathment considered here results in synaptic strengths and interaction kernels being drawn from entirely distinct distributions. This is a significant departure from many previous studies that consider heterogeneous connection strengths distributed according to a normal distribution, 𝒩 (*J, σ*) (Rajan and Abbott, 2006; Rajan et al., 2010; Mastrogiuseppe and Ostojic, 2018). Furthermore, this goes beyond the binary case (where each synapse was either ensheathed or unsheathed) considered in the spiking simulations in Handy and Borisyuk (2023), not only by considering multiple strengths at once but also through the development of this mean-field approximation. Lastly, this mean-field approximation runs at a fraction of the computational cost of spiking simulations, which require long runtimes for accurate estimation of the power spectrum and cross-spectrum.

### 3.3 Investigating the impact of glial ensheathment PV vs. SST interneurons

We now utilize the mean-field approximation to perform a broader parameter sweep and disentangle the effects of ensheathing both PV and SST interneurons. Specifically, we will consider networks with *either* PV or SST interneuron ensheathment across a range of ensheathment probabilities and strengths. To simplify this exploration, we consider only binary levels of ensheathment, where every synapse from population *b* is either ensheathed with strength *s*_*en*_ with probability *ρ*_*b*_ or unsheathed with probability 1 −*ρ*_*b*_.

Fig 7 shows the results for the ensheathment of PV neurons. We find that across a range of probabilities and strengths, this reliably leads to an increase in firing rates across the three populations, with larger deviations from baseline occurring for greater values of *s*_*en*_ and *ρ*_*p*_ (Fig 7a). Furthermore, we observe increases in the power spectrum of the excitatory population over a range of lower frequencies, along with a consistent increase in gamma power (Fig 7b and 7c). The coherence, specifically the gamma coherence, also increases for all values of *s*_*en*_ and *ρ*_*p*_ (Fig 7d and 7e).

**Fig 7.**
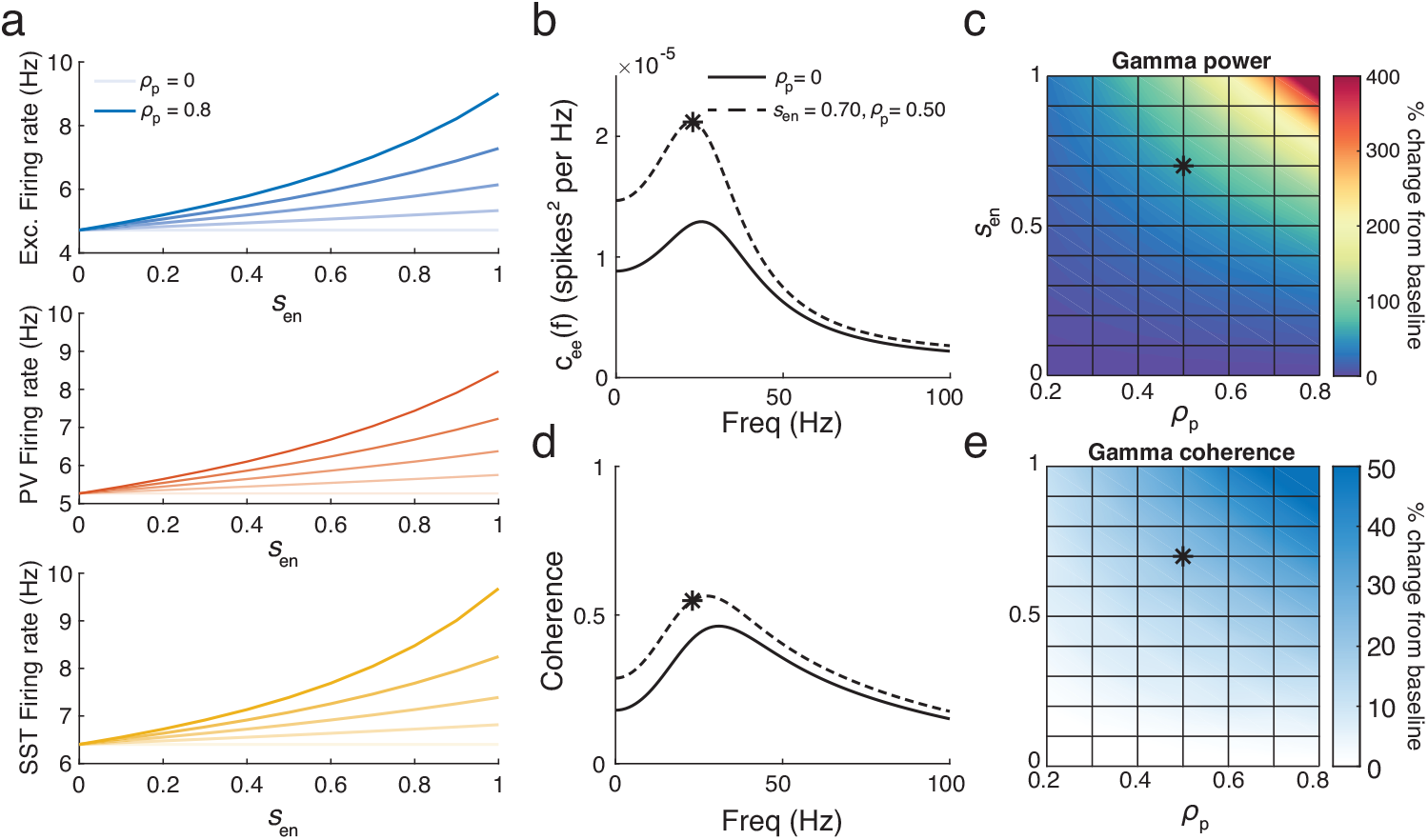
a: Firing rates of excitatory (top), PV (middle) and SST (bottom) neurons for a range of ensheathment strengths and probability of ensheathment of PV synapses (*ρ*_*p*_ = 0, 0.2, 0.4, 0.6, 0.8 from light to dark shades). b: Power spectrum for the excitatory population for *ρ*_*p*_ = 0 (solid) and 0.5 (dashed) for *s*_*en*_ = 0.7. The star indicates the gamma power. c: The percentage change in gamma power from baseline (*ρ*_*p*_ = 0) for a range of ensheathment strengths and probabilities *ρ*_*p*_. The star indicates the point in parameter space used in panel b. d: The same as panel b but for coherence. e: The same as panel c, but for gamma power

Fig 8 shows the results for the ensheathment of SST neurons. Similar to the previous findings, there is a reliable increase in firing rates across the three populations, indicating that the ensheathment of SST neurons is sufficient to drive the network into a hyperactive state (Fig 8a). However, the results for the power spectrum and coherence differ significantly (Fig 8b-e). The increases in the power spectrum are much more moderate, with gamma power not even doubling for the most extreme values of *s*_*en*_ and *ρ*_*s*_. Furthermore, the coherence changed only slightly as *s*_*en*_ and *ρ*_*s*_ increased. Overall, while the qualitative increases observed in Section 3.1 (particularly those found in Fig 6) could be achieved with the ensheathment of PV neurons alone, the corresponding increases in the power spectrum and coherence measures would not be replicated with the ensheathment of SST neurons alone.

**Fig 8.**
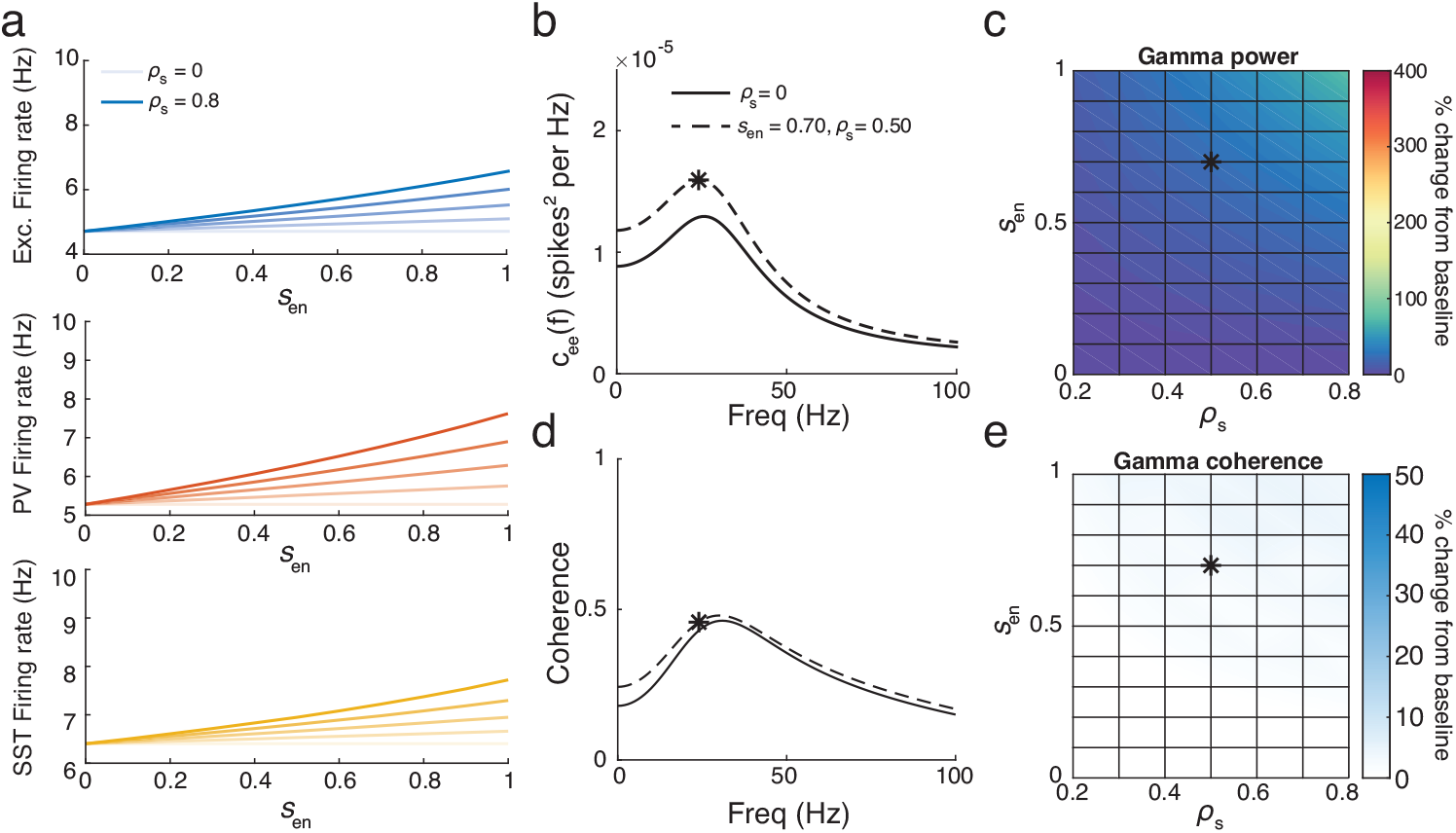
a: Firing rates of excitatory (top), PV (middle) and SST (bottom) neurons for a range of ensheathment strengths and probability of ensheathment of PV synapses (*ρ*_*p*_ = 0, 0.2, 0.4, 0.6, 0.8 from light to dark shades). b: Power spectrum for the excitatory population for *ρ*_*s*_ (solid) and 0.5 (dashed) for *s*_*en*_ = 0.7. The star indicates the gamma power. c: The percentage change in gamma power from baseline (*ρ*_*p*_ = 0) for a range of ensheathment strengths and values *ρ*_*s*_. d: The same as panel b but for coherence. e: The same as panel c, but for gamma power

## 4 Conclusion and discussion

As experimental techniques continue to advance and uncover additional heterogeneities within neuronal (and non-neuronal) sub-populations, simplified mean-field approximations will continue to be essential for exploring increasingly expansive parameter sets and highlighting key components behind complex processes. In this work, we present such an approximation derived from linear response theory that captures the effects of glial ensheathment on synapses. This theory successfully reproduces recent experimental findings, demonstrating that the network enters a hyperactive state when glial ensheathment of inhibitory synapses is present. It also enables us to examine not only average firing rates but also higher-order network statistics, leading to the testable prediction that both gamma power and gamma coherence significantly increase in the presence of glial ensheathment of inhibitory synapses. Further parameter exploration emphasized the importance of considering more than just average firing rates and reinforced the hypothesis of a division of labor across inhibitory subtypes (Bos et al., 2020a). While the ensheathment of PV *or* SST interneurons was sufficient to increase the network’s firing rate, only the ensheathment of PV interneurons led to significant increases in gamma power and gamma coherence.

Glial ensheathment is a dynamic process that remains under investigation. In this work, we focused on the findings of Haruwaka et al. (2024), which observed significant increases in microglial ensheathment of inhibitory synapses before and after anesthesia induced by the administration of isoflurane. This represents a substantial disruption to the system, as firing rates drop considerably in the anesthetized state. These results support the idea that microglia dynamically respond to the low-activity network state by attempting to boost excitatory firing rates, resulting in a rebound effect that manifests as hyperactivity during the emergence from anesthesia. Our theoretical prediction of increased gamma power and gamma coherence (along with a general increase across the power spectrum) during this emergence suggests additional, unintended consequences brought on by increased ensheathment. Since neuronal correlations play a key role in how the brain encodes information and performs computations (Cohen and Maunsell, 2009; Cohen and Kohn, 2011; Ruff and Cohen, 2014; Hazon et al., 2022; Kohn et al., 2016), it will be interesting to see if future experimental studies observe similar increases in glial ensheathment in other brain states with sustained aberrant firing rates, and whether such spurious correlations lead to cognitive impairments. If so, our prediction that PV interneurons are largely responsible for the observed increase in the power spectrum suggests a potential therapeutic avenue—specifically targeting those synapses to release them from glial ensheathment.

These results arise from only one possible effect of glial ensheathment, namely the shielding of the post-synaptic terminal via the clearance of neurotransmitters from the synaptic cleft. This effect is likely shared by two types of glial cells, microglia and astrocytes, and our model does not differentiate between the two. However, there are other pathways through which these cells can impact network dynamics, including synaptic pruning, adjustment of resting potentials via ion buffering, and the release of gliotransmitters (Paolicelli et al., 2011; Larsen et al., 2014; Covelo and Araque, 2018). The strength of these additional pathways will likely depend on the proximity of the glial cell to the synapse. Future work should aim to extend the microscale model presented here to account for these mechanisms and provide a more complete view of an “effective” glial cell. We anticipate that the mean-field approximation derived here can be extended to account for these additional pathways. For example, in the case of the reversal potential, once the effect of ensheathment is understood, the corresponding parameter *E*_*L*_ could be extended as a function of *s*_*en*_ and incorporated into the theory in a similar fashion to how we treated synaptic strength and time constants. Our mean-field approximation (Eq 15) relied on numerical computations to estimate the steady-state firing rate, the linear response function for the neuron subclasses, and the power spectrum of the unperturbed system. Analytical expressions for these functions have been derived for simplified leaky integrate-and-fire neurons, but they rely on non-intuitive mathematical functions like the complementary error function and parabolic cylinder functions (Holden, 1976; Lindner and Schimansky-Geier, 2001; Lindner et al., 2002). While we lack such analytical expressions, the exponential integrate-and-fire neuron used here offers the benefit of a more biologically realistic spiking mechanism, while remaining computationally efficient for our spiking neuronal network. We also left the mean-field approximation in matrix form, as its expansion would be largely uninterpretable. While concise formulas can be derived for simplified networks consisting of one or two populations of neurons (Lindner et al., 2005), this task becomes significantly more complex as the number of populations increases. Nevertheless, our theory enabled us to perform a large parameter sweep of the underlying system, revealing a non-intuitive result. Furthermore, it derived terms that deviate from a naïve mean-field approach, which could guide future mean-field approaches considering similar types of heterogeneity.

Overall, this work provides a foundation for understanding how the heterogeneities introduced by glial ensheathment influence network dynamics and offers a framework for future investigations that examine the role of glial cells in shaping complex neural computations.

## Data availability

The code used to generate the computational data and reproduce the figures of the paper will be deposited on Zenodo, an open research repository, upon paper’s acceptance.

## Acknowledgments

This work was supported by the Burroughs Wellcome Fund’s Career Award at the Scientific Interface to G. Handy.

